# The role of the unicellular bottleneck and organism size in mediating cooperation and conflict among cells at the onset of multicellularity

**DOI:** 10.1101/2023.07.17.549265

**Authors:** Sydney Ackermann, Matthew Osmond

**Affiliations:** Department of Ecology and Evolutionary Biology, University of Toronto

## Abstract

Evolutionary transitions in individuality introduce new levels of selection and thus enable discordant selection, threatening the stability of the transition. Cheating is such a problem for multicellularity. So why have so many transitions to multicellularity persisted? One possibility is that the unicellular propagule maintains cooperation among cells by purging cheaters. The evolution of propagule size has been modeled previously, but in the absence of competition between individuals, which may often select for larger propagules. How does the nature of competition between individuals affect the optimal propagule size in the presence of cheating? Here we take a model of early multicellularity, add phenotypic switching between cheating and cooperative phenotypes, and simulate size-dependent competition on a lattice, which allows us to tune the strength of interspecific vs. intraspecific competition via dispersal. As expected, cheating favors strategies with unicellular propagules while size-dependent competition favors strategies with few large propagules (binary fragmentation). How these opposing forces resolve depends on dispersal. Local dispersal, which intensifies intraspecific competition, favors binary fragmentation, which reduces intraspecific competition for space, with one unicellular propagule. Global dispersal instead favours multiple fission when cheating is common. We also find that selfishness promotes smaller body size, despite direct opposing selection from competition. Our results shed light on the evolution of multicellular life cycles and the prevalence of a unicellular stage in the multicellular life cycle across the tree of life.

**Author summary:** A multicellular organism is a group of cooperating cells. But wherever there is cooperation there is the temptation to cheat. Having offspring that start as a single cell (a unicellular bottleneck) has been hypothesized as an adaptation to purge lineages of cheating cells. We model the evolution of offspring size but add competition between individuals, which may select against small unicellular offspring. We find that having some unicellular offspring is still a successful strategy, but how many depends on the nature of competition.

## Introduction

Selection acts on variation. Where there is heritable variation across individuals – be it at the level of the gene, cell, organism, or beyond – there is potential for evolution by natural selection (Lewontin 1970). Selection acting at multiple levels simultaneously is known as multilevel selection (Szathmary and Maynard Smith 1995). Multilevel selection is a fundamental aspect of evolutionary transitions in individuality (Buss 1987) – such as the transitions from unicellular to multicellular life, from unlinked replicators to chromosomes, and from solitary organisms to eusociality (Szathmáry and Maynard Smith 1995) – as fitness is transferred to new units of selection (Folse and Roughgarden 2010, Hammerschmidt et al. 2014, Ratcliff et al. 2017, Howe et al. 2022).

A multicellular organism is composed of cooperating cells (Brunet and King 2017). This life-history strategy has independently evolved multiple times (Grosberg and Strathmann 2007, Niklas and Newman 2013, Atkipis et al. 2015), often resulting in evolutionary radiations (Herron et al. 2009). There are a number of potential advantages of multicellularity (Tong et al. 2022). One proposed advantage of multicellularity is the potential to be larger (Michod and Roze 2001). Indeed, multicellularity has evolved experimentally in response to selective pressures for larger size, either to avoid predation (Boraas et al. 1998) or to settle faster (Ratcliffe et al. 2012). Other possible advantages of multicellularity include, but are not limited to, division of labor, improved extracellular metabolism and cross-feeding (Ispolatov et al. 2012, Tong et al. 2022).

Conflict between levels of selection can arise when the units at these different levels each have influence over a phenotype with opposing effects on fitness (Lachmann et al. 2003). For example, in the case of multicellularity, cell replication rate can have very different effects on the fitness of the cell and the fitness of the organism. When mutations overwrite the checkpoints that maintain cooperation between cells, selection on replication rate at the cell level can result in cancer (Howe et al. 2022). Cancer (cheating cells) is a problem that all multicellular organisms must deal with and can be found across the tree of life, in vertebrates, plants, hydra, and mollusks to name a few (Atkipis et al. 2015). As much as cheating comes hand in hand with cooperation, cancer comes with multicellularity.

Most multicellular organisms begin as a unicellular propagule (Bell and Koufopanou 1991). In fact, the unicellular propagule has been described as a necessary condition for the evolution of obligate multicellularity (Howe et al. 2022). Why is it that multicellular organisms go to the trouble of starting from a single cell, temporarily forgoing all the benefits that come with multicellularity? One hypothesis is that the unicellular propagule is an organism-level adaptation to maintain cooperation among cells, i.e., to prevent cheating (Grosberg and Strathmann 2007, Grosberg and Strathmann 1998). The single-cell bottleneck purges lineages of cheating cells, preventing them from being vertically transmitted, by forcing selection on the cell and organism to align when the cell and the organism are one and the same.

A number of mathematical models shed light on why the unicellular propagule is such a common strategy. Building on previous work (Otto and Orive 1995; Michod 1996; Michod 1997), Roze and Michod (2001) asked what number of cells in the propagule maximizes population growth rate when cheating cells continually arise from mutation and the fitness of an organism depends on both the proportion of its cells that are cheating and its total number of cells (organism size). They found that the unicellular propagule, which reduces mutational load, maximizes population growth rate when the benefits of larger size are not too great. This model assumes non-overlapping generations, large adult sizes (since the cells in a propagule are sampled with replacement from the adult), and indeterminant growth.

A new approach to modeling the evolution of propagule size, more suitable for smaller, more primitive multicellular organisms, was introduced by Pichugin et al. (2017). Instead of choosing cells from an adult (with replacement) to create many propagules of the same size, they introduced the concept of fragmentation mode, defined by the maximum size of an organism and the sizes of the offspring it fragments into. The possible fragmentation modes for a given maximum size, *L*, are all partitions of *L*. As an example, consider *L*=4. This organism could fragment into all single cells (1+1+1+1), termed multiple fission; two groups of equal size (2+2), termed binary fission; one group of two and two single cells (2+1+1); or a group of three and one single cell (3+1), termed the unicellular propagule mode (Pichugin et al. 2017). To be clear, all fragmentation modes that include a single-celled offspring employ the more general phenomena of the unicellular bottleneck. Meanwhile the “unicellular propagule mode” defined by Pichugin et al. (2017) refers only to modes that fragment into just two groups, with one group being a single cell. This fragmentation framework captures the diversity of life-cycles we see in nature. For example, *Trichoplax adhaerens* reproduces via binary fission and bacterial biofilms reproduce via the unicellular propagule mode (Pichugin and Traulsen 2022; Ratcliff et al. 2017). Meanwhile, species in the genera *Chlamydomonas* and *Gonium* reproduce via multiple fission (Hanschen et al. 2016; Pichugin and Traulsen 2022), as do Acoelomorpha worms, sea stars and sea cucumbers (Howe et al. 2022, Mladenov 1996).

Ignoring mutation, Pichugin et al. (2017) found that all fragmentation modes that maximize population growth rate create only two offspring. This includes the unicellular propagule mode, which is optimal even in the absence of cheating when being large has outsized benefits. It may therefore be, then, that the single-cell bottleneck does not primarily evolve to prevent cheating. Rather, it may simply maximize population growth rate (see also Ratcliff et al. 2013). Adding frequency-dependent competition between organisms (Ress et al. 2022) does not affect the fact that all optimal modes have two offspring, but does allow for complications such as coexistence and bistability. Phenotypic switching between two cell types was added to the fragmentation framework by Gao et al. (2019), who used a payoff matrix to describe the interactions between the cells. Similar to Pichugin et al. (2017), they found that the optimal strategies were the unicellular propagule mode and binary fission (e.g., 2+2), but also multiple fission into all single cells (e.g., 1+1+1+1) when a group composed of both cell types is disadvantageous at the group level.

While some of the above models support the idea that the unicellular propagule can increase cooperation between cells in the face of cheating, for simplicity they do not include interactions both between cells and between organisms. However, given that cheating is a multilevel selection problem, it may be informative to consider interactions between individuals at multiple levels. In particular, here we ask, are unicellular propagules part of the optimal life-history strategy in the face of cheating when there is also size-dependent competition between organisms? Previous models of propagule evolution also ignore space. Because space interacts strongly with competition (Durrett and Levin 1994, Kerr et al. 2002, Boerlijst and Hogeweg 1991, Hermsen 2022), tuning the strength of interspecific vs. intraspecific interactions, here we also ask how the unicellular bottleneck fares with and without spatial structure.

In the presence of both cheating and size-dependent competition between organisms we expect that producing unicellular propagules will be part of the optimal strategy when the benefits of purging cheaters outweigh the costs of having small offspring. For example, as the phenotypic switching rate increases we expect that strategies that produce unicellular propagules will outcompete those that do not. To test this hypothesis we simulate the fragmentation framework of Pichugin et al. (2017) on a lattice, where each vertex can contain at most one group of cells, with recurrent switching between cooperative and cheater cellular phenotypes. We find that cheating, competition, and space together can select for many different life histories. In particular, increasing switching rates favors the production of unicellular propagules, as expected. However, in the absence of spatial structure, the optimal strategy at high switching rates is multiple fission, not the unicellular propagule strategy. Adding spatial structure alters this conclusion, favoring the unicellular propagule strategy.

## Methods

Below we first give an overview of our assumptions followed by a description of the full simulation method. The simulation code, written in Python, is publicly available on GitHub (https://github.com/Sydneya1/Lattice-Model).

We aim to model interactions among cells and groups of cells (individuals) at the origin of multicellularity. The organism we are modeling is hypothetical as we do not know exactly what primitive multicellularity looked like and our goal is to construct a simple, general model from which to glean qualitative results.

We assume cells divide clonally by mitosis and stick together after division to create a multicellular individual. Following Pichugin et al. (2017), we assume an individual has a fixed maximum cell number and once this cell count is reached it fragments according to a fixed partition (e.g., an individual grows to size 4 and fragments into individuals of size 1 and 3). The new individuals (offspring) inherit the fragmentation mode of their parent (corresponding to the pure life-cycle of Pichugin et al. 2017).

Following Gao et al. (2019), we assume a cell can have one of two phenotypes and can switch between the two upon division. Here we refer to the two cell types as cooperators and cheaters. We assume cheaters divide faster but increase the chances of group death (like cancer), with groups composed entirely of cheaters dying with certainty. This simple trade-off creates discordant multilevel selection that describes a key aspect of the evolutionary transition to multicellularity. Upon fragmenting, the cells of an individual are randomly assigned to the new groups, meaning that cheating cells are passed from parents to offspring (see Domazet-Loso and Tautz 2010 for an example of vertical transmission of cancer in hydra).

To simultaneously model competition between individuals, we place individuals on a lattice and allow at most a single individual at each vertex (see Figure 1). The use of a lattice also allows us to investigate the effect of spatial structure. Upon fragmenting, each offspring is assigned a vertex (either at random across the entire lattice – no spatial structure – or around the parent). If that vertex is already occupied the offspring competes with the resident to determine who survives. We model this competition as random (i.e., a fair coin flip) or with an advantage to larger individuals.

**Figure 1.**
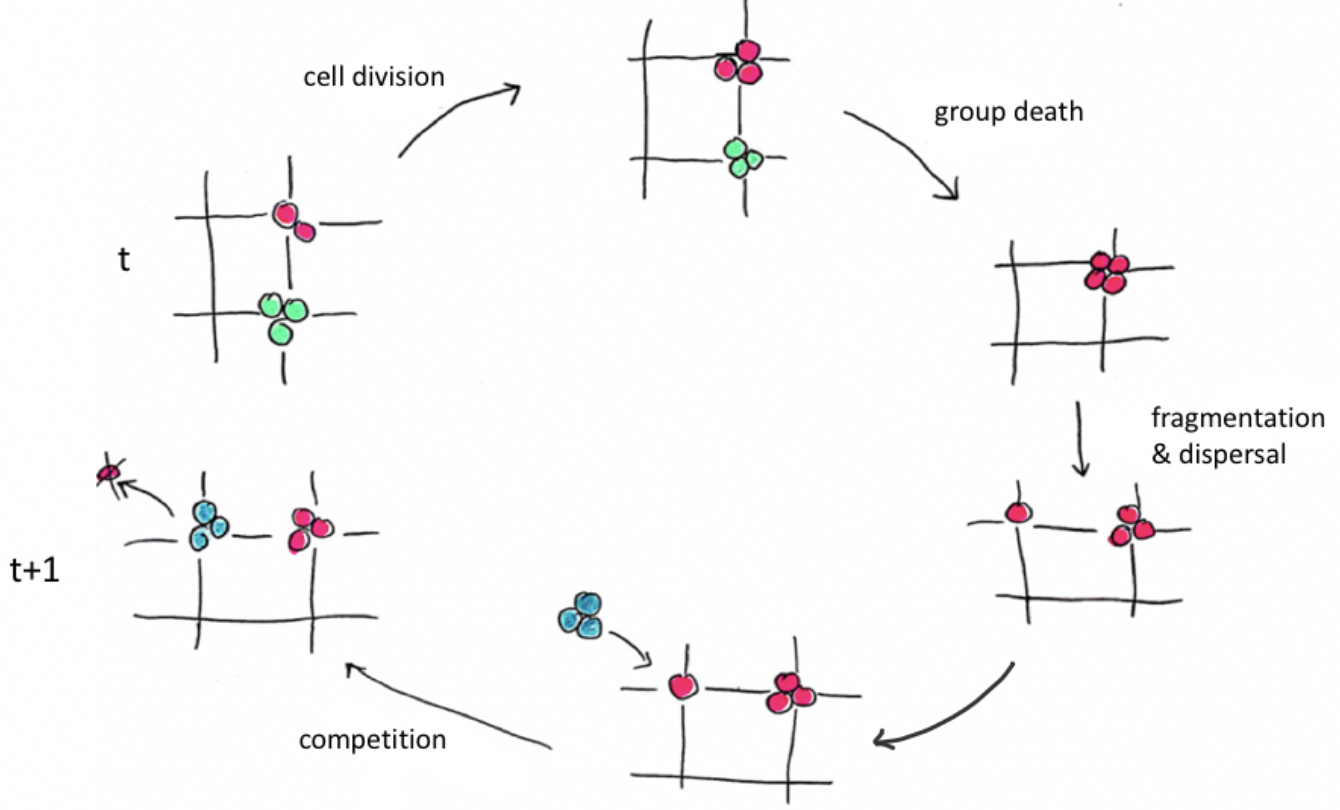
A sample of the lattice is shown to illustrate the simulated life cycle. The colours indicate the fragmentation modes, with pink being 3+1 (the modes of green and blue are not indicated here as they do not fragment). We start on the left with two groups, pink of size 2 and green of size 3. During the cell division stage the pink group grows by one cell, while the green group does not grow. The next stage is group death. In this instance the green group dies. Next the pink group fragments since it has reached its maximum size of 4. Both pink offspring are assigned to empty vertices, but elsewhere on the lattice an offspring of size three (with the blue fragmentation mode) is assigned to the spot that the pink single cell currently occupies. In this case the blue offspring wins the spot and the pink offspring dies (as would happen with complete size-dependent competition). After fragmentation the time is updated and the life cycle begins again. Phenotypic switching between cooperative and cheater cells (not shown) occurs during cell division.

Our interest is in which theoretically possible life-cycles dominate and, in particular, whether these dominant life-cycles produce unicellular offspring, which improve the ability of a lineage to purge cheaters (unicellular bottlenecks) but may be competitively inferior. To investigate this, we populate a lattice with a variety of fragmentation modes and simulate the dynamics forward in time. For simplicity we primarily simulate very small organisms (maximum 4 cells) but confirm that our key results continue to hold for larger sizes (maximum 10 cells). Note that while we model multilevel selection quite generally, here we focus on multicellularity for concreteness and because we are interested in cancer.

To start the simulation, *N*_*0*_ single cells are randomly scattered on an *M* x *M* lattice and each given a fragmentation mode. A fragmentation mode is defined by the maximum size of the group (adult size), *L*, and the way the adult fragments into offspring (a partition of *L*).

Each iteration starts with cell division (birth). Let the rate at which a cooperator cell divides in a group with *i* cooperator cells be *b*_*i*_. Following Pichugin et al. (2017), we define

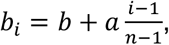

where *n* is the largest adult size among the modes allowed in a given simulation and *a* is the maximum additional birth rate, achieved when a group is maximal size and composed entirely of cooperators, *i*=*n*. Below we will consider two types of birth rates: 1) independent, in which the birth rate of a cell is independent of the group, *a*=0, and 2) cooperator-dependent, in which the birth rate of a cell increases linearly with the number of cooperators in the group, *a*=1. The probability a given cooperator cell divides in a group with *i* cooperators is then

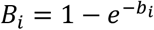

and the number of cooperator cells that divide is a binomial random variable.

The division rate of a cheater cell in a group with *i* cooperators is

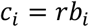

where *r* >1 is the replicative advantage of cheaters. The probability a cheater divides in a group with *i* cooperators is then

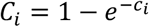

and the number of cheater cells that divide is a binomial random variable. We set *b*_*i*_ = *c*_*i*_ = 0 so that cheaters can only divide if a cooperator cell is present.

Note that if cell division results in the adult size being exceeded then newly formed cheater and cooperator cells are randomly chosen until the adult size is reached. When a cooperator cell divides, the new cell has probability *m* of switching to a cheater cell.

Similarly, when a cheater cell divides the new cell switches to a cooperator cell with probability w.

The next event in a time step is group death. There is a constant background mortality rate, *d*. This death rate is converted to a probability

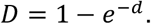

Groups also die with probability *P*, where *P* is the proportion of cheaters in the group. This implies that groups consisting entirely of cheater cells, including a solitary cheater cell, always die. Groups survive both forms of mortality with probability (1-*D*)(1-*P*).

Finally, fragmentation, dispersal, and competition occur. Working through one group at a time, if it has reached its adult size it splits apart into offspring whose size and number are determined by its fragmentation mode. In this way, any cheater cells are randomly distributed among offspring. Each offspring is then randomly assigned a place on the lattice. Here there are two treatments, 1) global dispersal (no spatial structure) and 2) local dispersal (spatial structure). Under global dispersal each offspring has an equal chance of landing on any vertex on the lattice. Under local dispersal each offspring has an equal chance of dispersing to one of five neighboring vertices: on the parent vertex or one vertex above, below, right, or left of it.

In either case, if an offspring lands on a vertex that is already occupied, competition occurs. Under complete size-dependent competition the larger group will win the spot (if the vertex is occupied by a group of the same size then each has a 50% chance of winning the spot). Individuals that lose the spot die. To isolate the effects of size dependence, we also model random competition, where every offspring has a 50% chance of winning an occupied vertex regardless of size. The gradient in-between these two extreme forms of competition can be explored by introducing a parameter *s* such that the probability of the larger individual winning a spot is (1+*s*)/2, with *s* = 0 corresponding to random competition and *s* = 1 corresponding to complete size-dependent competition.

This process repeats for *T* timesteps. A visualization of this life cycle of cell division, group death, fragmentation, dispersal, and competition can be seen in Figure 1. An example of the lattice before and after the simulation, as well as a time series plot showing the dynamics of the population during the simulation can be seen in Figure 2. The parameters used in the simulation are summarized in Table 1.

**Table 1.** A description of the parameters used in the simulation.

| Parameter | Description | Default value (if applicable) |
| --- | --- | --- |
| $M$ | Length of the square lattice | 50 |
| $N_0$ | Initial number of groups on the lattice | |
| $L$ | Size at fragmentation, i.e., adult size | |
| $b_i$ | Division (birth) rate of cooperator cells in a group with $i$ cooperators | |
| $b$ | Baseline birth rate | 0.3 |
| $a$ | Maximum additional birth rate due to cooperation | 0,1 |
| $n$ | Maximum adult size in a given simulation | 4 |
| $B_i$ | Probability a cooperator cell in a group with $i$ cooperator cells divides | |
| $c_i$ | Division (birth) rate of cheater cells in a group with $i$ cooperators | |
| $r$ | Replicative advantage of cheater cells | 2 |
| $C_i$ | Probability a cheater cell in a group with $i$ cooperator cells divides | |
| $m$ | Probability a new cooperator cell switches to a cheater cell | 0.2 |
| $w$ | Probability a new cheater cell switches to a cooperator cell | m |
| $d$ | Rate of background group mortality | 0.03 |
| $D$ | Probability a group dies due to background mortality | |
| $P$ | Proportion of cheater cells in a group = the probability a group dies due to cheaters | |
| $s$ | The degree to which competition between groups depends on size | 0,1 |
| $T$ | Number of iterations in the simulation | |

**Figure 2.**
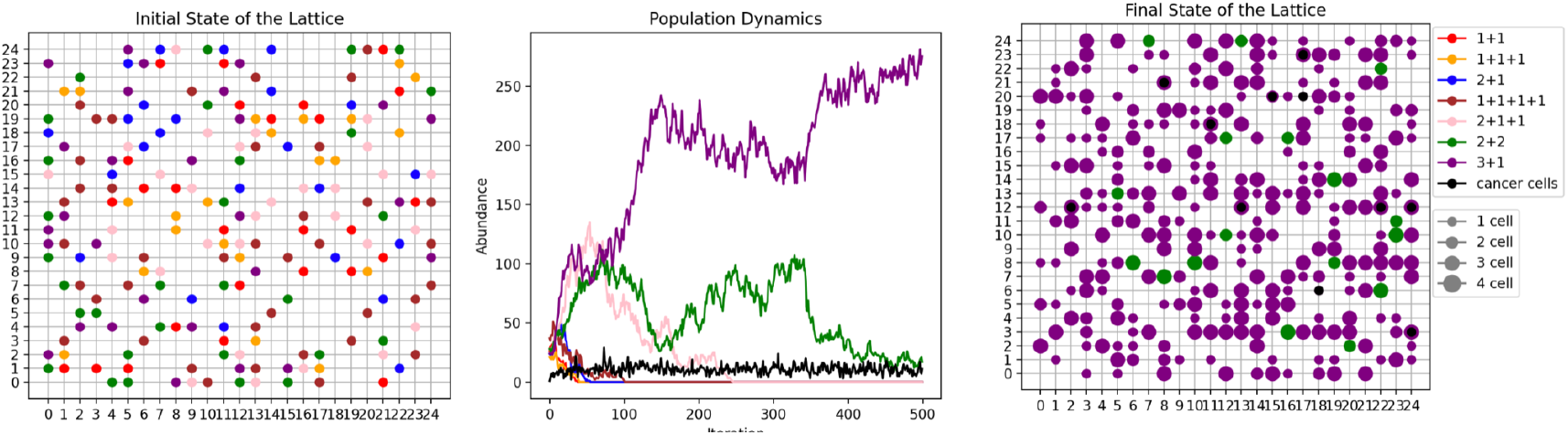
The initial state of the lattice, the abundances of each fragmentation mode (and the number of cheating cells) throughout the simulation, and the lattice after 500 timesteps. Parameters: *T*=500, *M*=25, *N*_*0*_=200, *n*=4, *b*=0.3, *d*=0.03, *r*=2, *m*=w=0.1, *s*=1, *a*=0, global dispersal.

## Results

### Dominant modes in the absence of cheating

Our model explores a combination of features that has not been considered previously. To isolate the effect of each feature we first ignore cheaters and phenotype switching (*m=w*=0) and focus on the effect of competition under different modes of dispersal. To do this we run simulations with either random competition, where a fragment has a 50% chance of winning an occupied spot regardless of size (*s*=0), or size dependent competition, where the larger individual always wins the spot (*s*=1), and either global or local dispersal. For each combination of competition and dispersal we run simulations across a range of birth rates with a maximum adult size *n*=4 and record the modes with the highest mean abundance (Figure 3).

**Figure 3.**
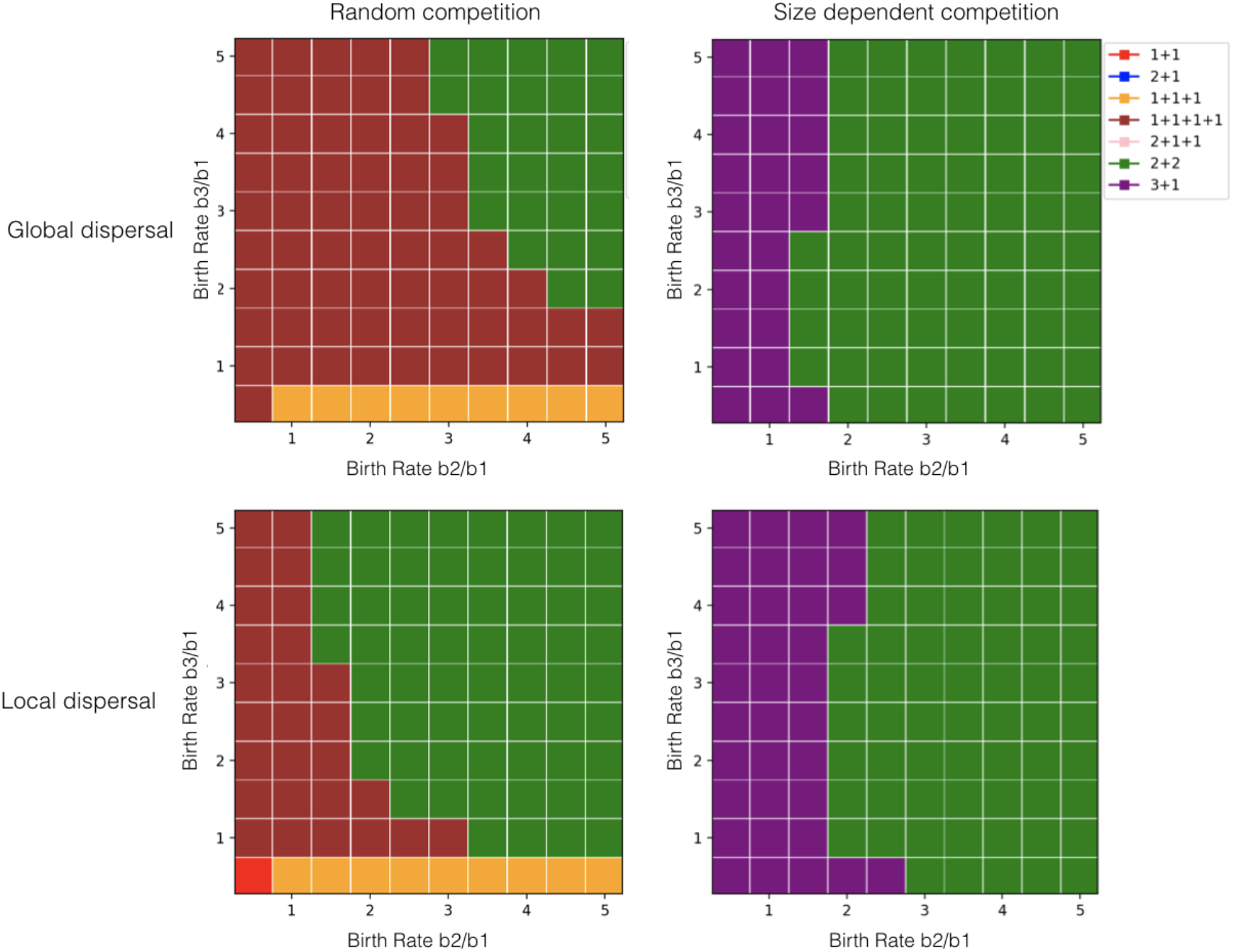
Dominant fragmentation modes in the absence of cheating (*m*=*w*=0). The dominant mode is the mode with the highest average abundance over 10 replicate simulations. Parameters: *T*=1000, *M*=50, *N*_*0*_=2000, *n*=4, *b*_*1*_=0.1, *d*=0, *s*=0 (left) and *s*=1 (right).

The top left panel of Figure 3 shows that the combination of random competition and global dispersal favors multiple fission (1+1+1+1) for most of the parameter space, except for when cells divide much faster in groups, where binary fission (2+2) wins, and when cells divide very slowly in groups of size three, where a smaller adult size is favored (1+1+1). This is in strong contrast to the outcome in the absence of competition, where the mode with the highest growth rate always fragments in two (see Figure 3A of Pichugin et al. 2017), and to the outcome with Lotka-Volterra-like size-independent competition, which also favors modes with exactly two offspring (Ress et al. 2022).

Here, under spatially-explicit size-independent competition, there is an advantage to fragmenting into more than two offspring. We believe the advantage of having multiple offspring arises from the fact that competition affects dispersing offspring and resident groups differently, which is not the case in Ress et al. (2022). The probability that a dispersing offspring gets a spot on the lattice is the probability it lands on an empty spot, 1 − *N*/*M*^9^, where *N* is the number of individuals on the lattice, plus the probability it lands on an occupied spot but wins, *N*/(2*M*^9^), which together give *p* = 1 − *N*/(2*M*^9^).

Meanwhile the probability that a resident group is not displaced by one particular offspring is the probability that the offspring does not land on it, 1 − 1/*M*, plus the probability that the offspring does land on it but fails to win, 1/(2*M*). Summing these and multiplying over all offspring produced in a given time step, O, gives *q* = (1 − 1/(2*M*))^*0*^. Since in general *p* ≠ *q*, the way a mode grows in number will impact its growth rate. To see this, imagine that *p* and *q* are constants and compare two modes, mode A where two offspring are produced every two time steps and mode B where four offspring are produced every four time steps. The number of groups of each mode after a large number of time steps, *t*, is expected to be *N*_*A*_(*t*) = ((1 − *D*)(*p*^3^ 4*q*))^*t*/2^ and *N*_*B*_(*t*) = ((1 −*D*)(*p*2*q*))^*t/*4^. If *p* = *q* then *N*_*A*_(*t*) = *N*_*B*_(*t*), and we have *N*_*A*_(*t*) < *N*_*B*_(*t*) whenever *q*<*p*, and vice-versa. In words, there is an advantage to modes that grow by having more offspring less often, over those that have less offspring more often, when the probability that a dispersing offspring survives competition is less than the probability that a resident group survives competition (*q*<*p*). We believe this explains the advantage of multiple fission over binary fission in our simulations. This advantage of having more offspring less often also allows multicellularity to dominate unicellularity when cells divide slower in groups (*b*_*2*_ < *b*_*1*_ and *b*_*3*_ < *b*_*1*_).

The bottom left panel of Figure 3 shows that restricting the dispersal of offspring reduces the advantage of multiple fission, as it dominates for less of the parameter space. Under local dispersal each mode competes more with itself, which is particularly damaging to a mode that has many offspring at one time, as this increases the chances that multiple offspring compete for the same spot on the lattice.

On the other hand, the right column of Figure 3 shows that size-dependent competition favors two binary fragmentation strategies: the unicellular propagule mode (3+1) and binary fission (2+2). When cells in groups of size 2 divide slowly (small *b*_*2*_ values), 3+1 wins because offspring of size 3 do not have to go through this slow stage. For larger *b*_*2*_ values 2+2 wins because groups of size 2 grow faster than groups of size 1. These results closely match those of Pichugin et al. (2017) (compare the top right panel of Figure 3 with their figure 3A) despite the fact that they do not model competition. The key difference is that Pichugin et al. (2017) see modes with smaller adult sizes (1+1 and 2+1) win when large groups grow slowly (*b*_*3*_ < *b*_*1*_). It therefore appears that size-dependent competition restricts the parameter space over which modes with smaller adult size win.

Under size-dependent competition, local dispersal favors the 3+1 mode over more parameter space (compare right panels in Figure 3). This likely results from the fact that the 3+1 mode has two very different offspring sizes. Competition within each mode (intraspecific competition) will favor the larger offspring, which will displace the smaller. This means that the 3+1 mode loses fewer cells to intraspecific competition than the 2+2 mode does, giving it a growth advantage. It also means that the average size of surviving offspring from the 3+1 mode will be larger than 2, giving it an advantage in interspecific competition as well.

A peculiar observation from these simulations is that the dominant modes under size-dependent competition (right panel of Figure 3) are more similar to the dominant modes in the absence of competition (Pichugin et al. 2017 figure 3A) – e.g., exclusively binary fragmentation – than they are to the dominant modes under random competition (left panel of Figure 3), which includes multiple fission. This may be explained by the fact that the two modes with the highest growth rate in the absence of competition over most of the parameter space (2+2 and 3+1) also have the largest possible adult (4) and mean offspring (2) sizes and therefore also do well under size-dependent competition.

### Dominant modes in the presence of cheating

After isolating the effect of size-dependent competition and its interaction with dispersal, we now combine this with switching between cooperative and cheating cell phenotypes. Instead of varying birth rates continuously we choose two different scenarios (independent birth, *a=0*, and cooperator-dependent birth, *a*=1) and vary the rate at which cells switch phenotype (*m,w*). For each parameter set we record the abundance of all modes at the end of the simulation and average over replicates (Figure 4).

**Figure 4.**
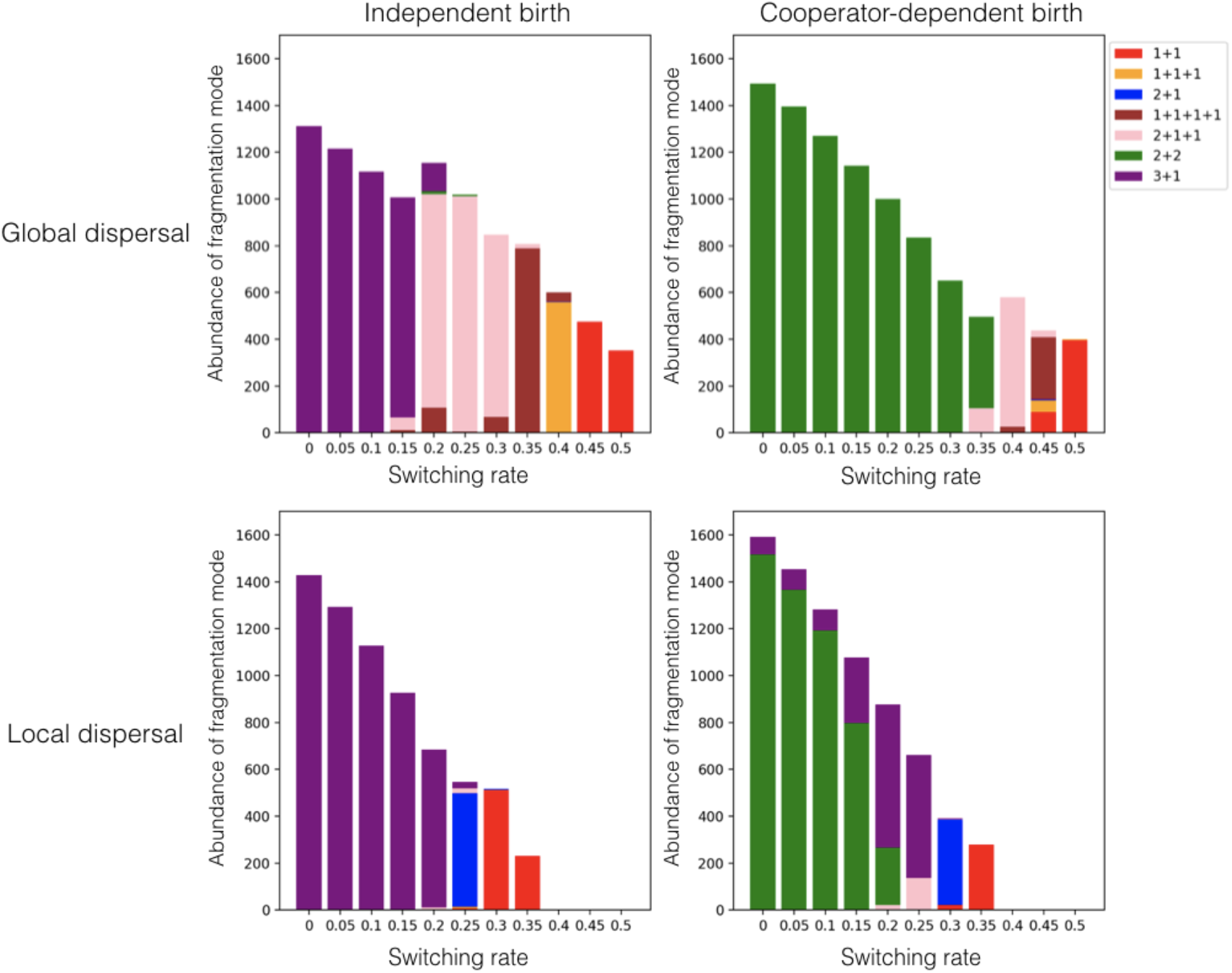
The final abundance of each fragmentation mode as a function of the switching rate. Abundances are averaged over 100 replicates. Parameters: *T*=5000, *M*=50, *N*_*0*_=2000, *n*=4, *b*=0.3, *d*=0.03, *r*=2, *s*=1, w=m.

Under independent birth and global dispersal (top left panel of Figure 4) we see the 3+1 mode winning in the cases of little to no phenotypic switching, which matches our earlier result (*b*_*2*_/*b*_*1*_=*b*_*3*_/*b*_*1*_=1 in the top right panel of Figure 3). As we increase the switching rate we see a progression of the most abundant modes from 3+1, 2+1+1, 1+1+1+1, 1+1+1, to 1+1. This progression shows that modes with a higher proportion of single celled offspring are selected for by cheating. In particular, multiple fission is the best strategy for high switching rates. This is consistent with the literature: unicellular bottlenecks purge lineages of cheating cells by aligning the fitness interests of the cell and the group. At very high switching rates we see a decline in the optimal adult size. This decline in adult size may be because individuals are too likely to die from cheating before fragmenting.

Next we look at cooperator-dependent birth rates and global dispersal (top right panel of Figure 4). Now cells in larger groups have a higher probability of dividing, e.g., because cells in a group can absorb and store resources more efficiently (Kaiser 2001). Binary fission (2+2) is the winning strategy with little or no cheating, consistent with previous results (*b*_*2*_/*b*_*1*_≅2 and *b*_*3*_/*b*_*1*_ ≅3 in the top right panel of Figure 3). As we increase the switching rate we see a progression in the most abundant mode from 2+2, 2+1+1, 2+1, 1+1+1, to 1+1. As under independent birth, this is an increase in the number of unicellular offspring and eventually a decline in adult size. However, because solitary cells divide slowest under cooperator-dependent birth, we do not see some multiple fission modes (1+1+1+1) and instead the larger-offspring binary fragmentation mode (2+2) dominates at relatively high phenotypic switching rates.

Local dispersal, and the resulting increase in intraspecific competition, increases the advantage of the unicellular propagule mode over multiple fission under both birth rate scenarios (bottom panels of Figure 4). In fact, only the unicellular propagule mode is observed under independent birth (bottom left panel of Figure 4). The increased intraspecific competition also causes a more precipitous drop in total population size and the extinction of all modes at lower mutation rates.

### Cheater load

To understand why particular modes dominate in the presence of cheating we run simulations with one mode on the lattice at a time and calculate the proportion of groups on the lattice that contain cheating cells (Figure 5).

**Figure 5.**
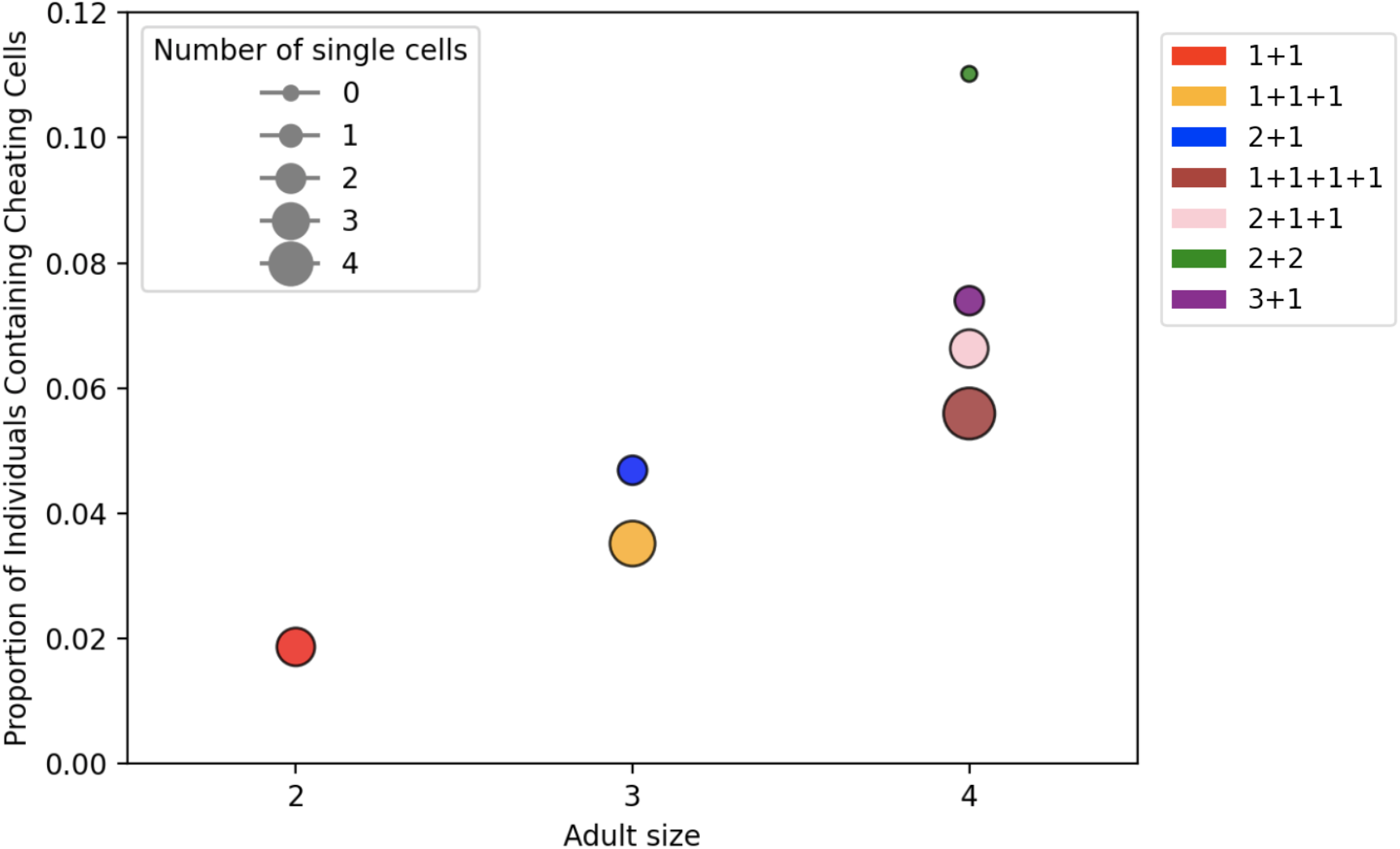
Cheater load as a function of adult size. The size of a circle indicates the number of single-cell offspring in that fragmentation mode. Parameters: *T*=1000, *M*=50, *N*_*0*_=1000, *n*=4, *b*=0.3, *d*=0.03, *r*=2, *m*=w=0.2, *s*=1, *a*=0, global dispersal, 80 replicates.

In Figure 5, for each adult size we see that multiple fission (1+1+…+1) has the lowest amount of cheating and binary fission has the highest. Moreover, the number of unicellular offspring does a very good job at ranking the cheater load of each mode. This sheds light on the patterns in Figure 4 – modes with more unicellular offspring have less cheaters in their population because solitary cheater cells die with certainty, purging cheating lineages. We also see an increase in the cheater load with adult size. Increasing adult size increases the number of cellular generations per organismal generation. This increases the number of chances a cheater cell arises as well as the strength of selection among cells relative to selection among organisms – resulting in larger loads in larger modes for a given number of unicellular offspring.

## Discussion

### Summary

By simulating competition between both cells and organisms in a spatial context we shed light on the evolution of early multicellular life cycles. First, we found that in the absence of cheating, independent (“random”) competition favors multiple fission, which minimizes the number of risky offspring establishment events needed to create a given number of groups, while size-dependent competition favors binary fragmentation, the strategies with the largest average offspring (Figure 3). Second, the effect of cheating depends on the dispersal mode (Figure 4). Under global dispersal, increasing the rate of switching to a cheating cell phenotype first favors modes with more unicellular offspring (such as multiple fission), then modes with smaller adult sizes. Under local dispersal, the unicellular propagule mode (binary fragmentation with one single-celled offspring) is favored at all rates of cancer and increasing the rate only decreases adult size.

Therefore, whether cheating can be said to explain the preponderance of species using the unicellular propagule mode will depend on the nature of dispersal. Third, we explored the mechanism behind these trends and found that the number of unicellular offspring is negatively correlated with a population’s cheater burden (Figure 5), confirming the ability of the unicellular bottleneck to purge cheaters.

### Major results

One of our key new results is the dominance of life cycles other than binary fragmentation. Previous models have found binary fragmentation to dominate whether considering discrete or continuous time (Pichugin and Traulsen, 2022), exponential or frequency-dependent growth (Pichugin et al. 2017, Ress et al. 2022), or even when considering bidirectional switching between two cell types (Gao et al. 2019). Here, however, we find modes that produce multiple offspring to dominate under a wide range of conditions. For one, by explicitly modeling competition in space, we see that multiple fission can dominate when single cells are not disadvantaged too much in growth or competition relative to larger groups (Figure 3). The reason for this is because a group performing multiple fission can produce a given number of replicates of itself with the minimum number of offspring. For example, it takes 4 offspring for the 1+1+1+1 mode to reach 4 groups but it takes 6 offspring for the 2+2 mode to reach 4 groups. Multiple fission is then favoured when the establishment of offspring is less likely than the persistence of resident groups. Adding cheaters can also cause multiple fission to dominate (Figure 4). In this case the reason is because unicellular offspring are an effective way for an individual to purge cheating from its descendants (Figure 5). One of the few factors previously found to promote multiple fission was costly fragmentation, where one cell is chosen to die at fragmentation (Pichugin et al. 2017). Curiously, this, like random competition and cheating, involves cell death. The other factor previously shown to favour multiple fission is a cost to groups of heterogeneous cells (Gao et al. 2019), which has some similarities with our model of cheating.

By simulating both global and local dispersal we identified the effect of spatial structure and its interaction with cheating. In the absence of cheating, local dispersal increased the parameter space over which the unicellular propagule mode dominated binary fission (right panel of Figure 3). We reason this to be due to increasing the relative strength of intraspecific (vs. interspecific) competition. Given competition is size dependent, intraspecific competition will disfavour smaller offspring. This means that the unicellular propagule mode will mostly lose unicellular offspring, which have a lower reproductive value than the larger offspring. Since the two offspring under binary fission are equivalent, intraspecific competition causes a larger loss of reproductive value in this mode. With cheating, local dispersal largely prevents modes with more than two offspring from dominating (Figure 4). Our reasoning for this is similar: having more offspring means dividing reproductive value into more, and hence more equal, pieces. Given intraspecific competition for space, more offspring means more death, and therefore a larger loss of reproductive value. By preventing multiple fission from dominating and, more generally, by decreasing the relative fitness of unicellular offspring, local dispersal hinders the purging of cheaters.

Throughout much of parameter space, increasing rates of cheating favours multiple fission (Figure 4) as it is the most efficient mode at purging cheaters (Figure 5). This supports our hypothesis that cheating will select for life cycles with unicellular offspring. On the other hand, it also forces us to clarify what we mean when we say that cheating selects for ‘the unicellular strategy’. It is often implied that humans, like many metazoans, partake in the unicellular strategy, since all offspring start from a unicellular zygote. However, in a model of overlapping generations like ours (and ignoring the complications of sex), humans would use something like the unicellular propagule mode with a very large adult size: reproduction creates one unicellular offspring and one surviving adult. More generally, the unicellular bottleneck is often synonymous with the unicellular propagule mode (Grosberg and Strathmann 2007, Grosberg and Strathmann 1998, Marquez-Zacarias et al. 2021, Avise 1993). Here we have shown that, while the unicellular propagule mode is more effective at minimizing cheater burden than some other modes, such as binary fission, multiple fission is the most effective. Of course, the costs and benefits of the unicellular bottleneck are experienced by any life cycle that has at least one unicellular offspring. We therefore suggest that the unicellular strategy be thought of as a continuum, with some modes using more of this strategy than others, with multiple fission at one extreme. As we have shown, the optimal amount of the unicellular strategy to use will depend on a variety of factors, such as the nature of competition and the rate of switching to cheater cells. Finally, we note that many previous models have assumed non-overlapping generations (e.g., Roze and Michod 2001), in which case cheating selects for a large adult that has many unicellular offspring, reminiscent of multiple fission.

In addition to more unicellular offspring, we also see a decrease in adult size in response to more cheating (Figure 4). This is because growing to a larger size requires more cell divisions, such that modes with a larger adult size have an increased risk of acquiring and dying from cheating cells before fragmenting. This risk is amplified by the fact that more cell divisions means an increased strength of selection among cells, further increasing the frequency of cheaters. The increased risk of dying from cheating can be seen in Figure 5, where groups with larger adult sizes are more likely to contain cheating cells. In the special case of cancer this implies that, all else being equal, cancer should select for smaller body sizes and larger individuals should have more cancer (Kokko and Hochberg 2015). But of course not all else is equal, and large organisms have found other ways to suppress cancer, leading to Peto’s paradox (Kokko and Hochberg 2015). However, at the onset of multicellularity, when the problem of cheating is initially introduced, other forms of cheater suppression are unlikely to have occurred yet, so that selection may initially act on any variation that exists in adult size.

### Future directions

One major limitation of our model is that the number of fragmentation modes grows rapidly with adult size, making it more complicated to explore what happens with larger organisms. For example, the family Volvocaceae includes organisms that are comprised of as little as 16 cells, but there are already 898 modes with an adult size of 16 or less. However, to check if the qualitative patterns we see with maximum adult size 4 hold more generally, we also performed simulations with maximum adult size 10 (giving 128 modes). This confirmed the general pattern of more single-celled offspring and smaller adult sizes with increasing switching rate (Figure S1), driven by a lower cheater load for modes with more single-celled offspring and smaller adult sizes (Figure S2). One important caveat to keep in mind when scaling our model up to larger sizes is that under local dispersal we assume there are only 5 vertices that the offspring can land on, which selects against modes with more offspring.

Here we competed modes against one another to determine the dominant strategies. A different approach would be to allow modes to mutate between each other and look for evolutionary stable strategies. The results of these two approaches could differ if there is strong frequency-dependence between modes. However, given that we do not see much variance in the dominant strategies under local dispersal between replicates, we do not expect the alternative mutation approach to change our qualitative results, at least for the relatively small organisms explored here.

## Acknowledgements

We thank Puneeth Deraje, Megan Frederickson, Nicole Mideo, Yuriy Pichugin, Locke Rowe, and Stephen Wright for their questions and comments. This work was supported by the Natural Sciences and Engineering Research Council of Canada (RGPIN–2021-03207 and DGECR-2021-00114 to MO). Computations were performed on the Niagara supercomputer at the SciNet HPC Consortium. SciNet is funded by: the Canada Foundation for Innovation; the Government of Ontario; Ontario Research Fund - Research Excellence; and the University of Toronto.

## Supplementary Figures Legend

Figure S1. The abundance of each fragmentation mode as a function of the switching rate for adult sizes up to 10 cells.

Figure S2. Cheater load as a function of adult size for adult sizes up to 10 cells.

