## Supplementary figures for "The role of the unicellular bottleneck and organism size in mediating cooperation and conflict among cells at the onset of multicellularity"

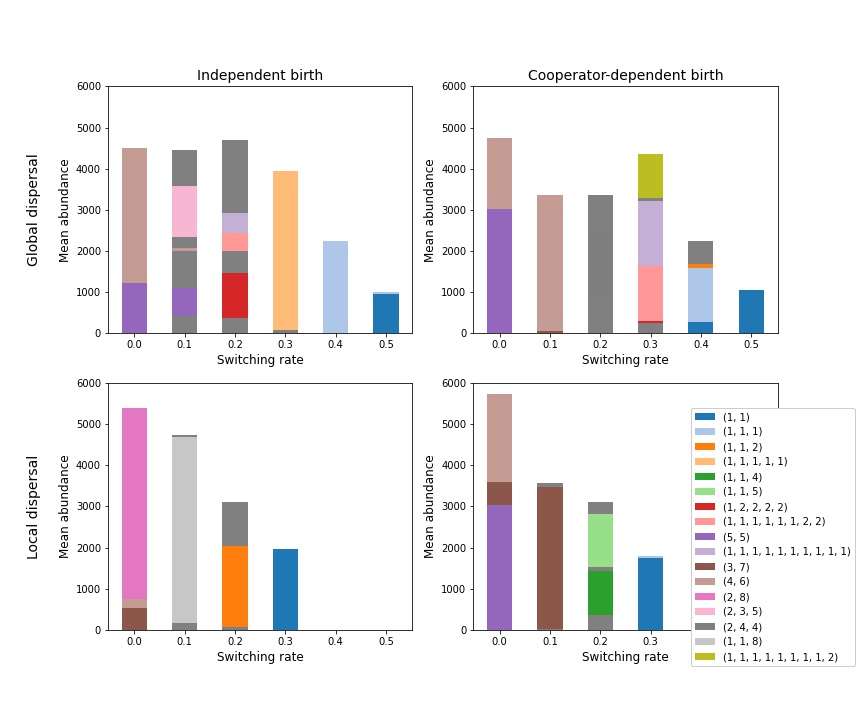

**Figure S1.** The abundance of each fragmentation mode as a function of the switching rate. Abundances are averaged over 10 replicates. For simplicity, only those modes that have a mean abundance greater than 1000 at any one switching rate are colored and shown in the legend. Parameters: *T*=5000, *M*=100, *N_0_*=10000, *n*=10, *b*=0.3, *d*=0.03, *r*=2, *s*=1, w=m. Compare to Figure 4, where n=4.

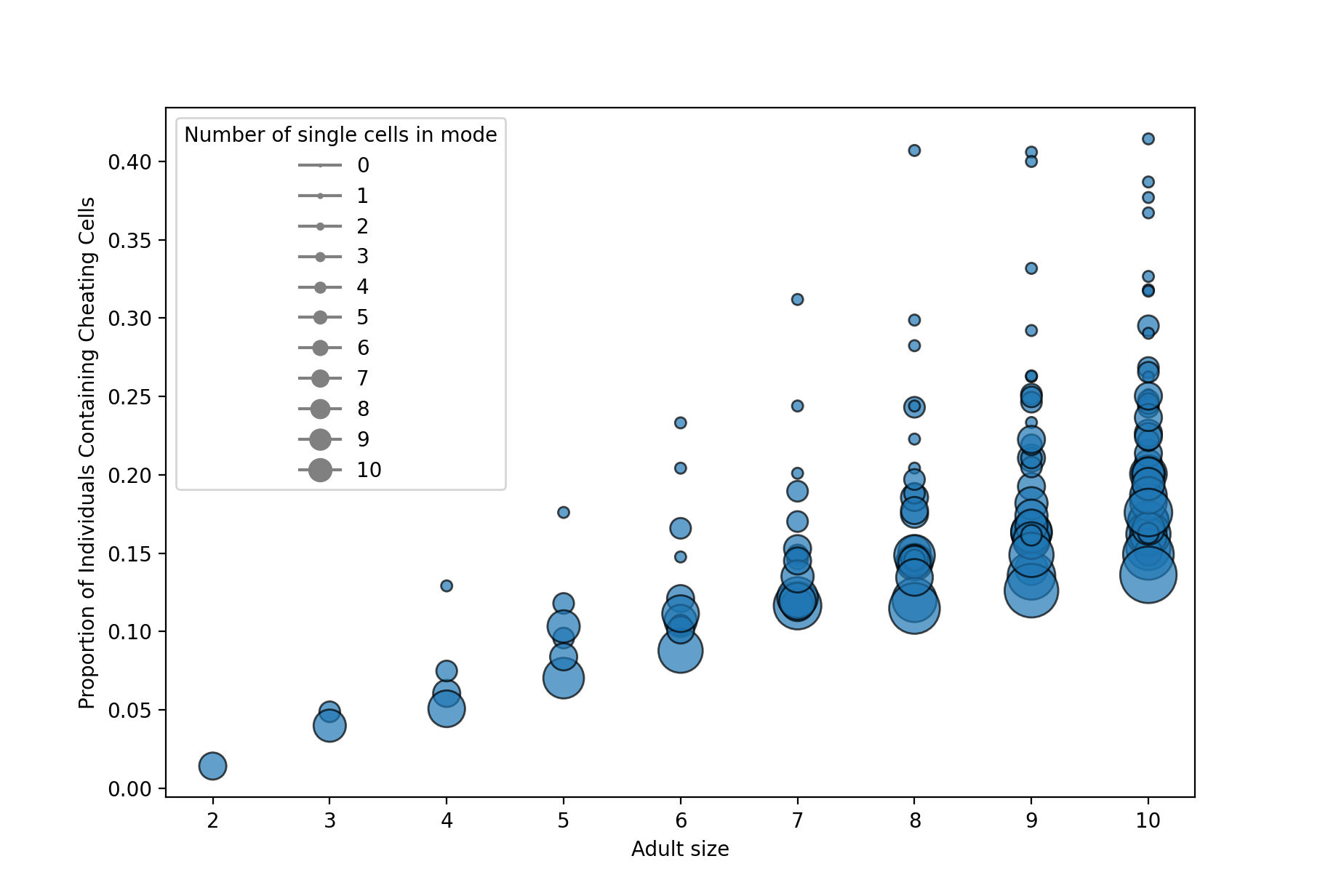

**Figure S2.** Cheater load as a function of adult size. The size of a circle indicates the number of single-cell offspring in that fragmentation mode. Parameters: *T*=1000, *M*=50, *N_0_*=1000, *n*=10, *b*=0.3, *d*=0.03, *r*=2, *m*=w=0.2, *s*=1, *a*=0, global dispersal, 1 replicate. Compare to Figure 5, where n=4.
